# Characterization of Novel Human β-glucocerebrosidase Antibodies for Parkinson Disease Research

**DOI:** 10.1101/2023.09.14.557851

**Authors:** Tiffany Jong, Alexandra Gehrlein, Ellen Sidransky, Ravi Jagasia, Yu Chen

## Abstract

**BACKGROUND:** Mutations in *GBA1*, which encodes the lysosome enzyme β-glucocerebrosidase (also referred to as acid β-glucosidase or GCase), are the most common genetic risk factor for Parkinson disease (PD) and dementia with Lewy bodies (DLB). Evidence also suggests that loss of GCase activity is implicated in PD without *GBA1* mutations. Consequently, therapies targeting GCase are actively being pursued as potential strategies to modify the progression of PD and related synucleinopathies. Despite this significant interest in GCase as a therapeutic target, the lack of well-characterized GCase antibodies continues to impede progress in the development of GCase-targeted therapies.

**OBJECTIVE:** This study aims to independently evaluate human GCase (hGCase) antibodies to provide recommendations for western blot, immunofluorescence, immunoprecipitation, and AlphaLISA (Amplified Luminescent Proximity Homogeneous Assay) assays.

**METHODS:** Two mouse monoclonal antibodies, hGCase-1/17 and hGCase-1/23, were raised against hGCase using imiglucerase, the recombinant enzyme used to treat patients, as the antigen. These novel antibodies, alongside commonly used antibodies in the field, underwent evaluation in a variety of assays.

**RESULTS:** The characterization of hGCase-1/17 and hGCase-1/23 using genetic models including *GBA1* loss-of-function human neuroglioma H4 line and neurons differentiated from human embryonic stem cells (hESCs) revealed their remarkable specificity and potency in immunofluorescence and immunoprecipitation assays. Furthermore, a hGCase AlphaLISA assay with excellent sensitivity, a broad dynamic range, and suitability for high throughput applications was developed using hGCase-1/17 and hGCase-1/23, which enabled a sandwich assay configuration.

CONCLUSIONS

The hGCase immunofluorescence, immunoprecipitation, and AlphaLISA assays utilizing hGCase-1/17 and hGCase-1/23 will not only facilitate improved investigations of hGCase biology, but can also serve as tools to assess the distribution and effectiveness of GCase-targeted therapies for PD and related synucleinopathies.

## Introduction

The lysosomal enzyme glucocerebrosidase (GCase) (EC 3.2.1.45), encoded by the gene *GBA1*, hydrolyzes its lipid substrate glucosylceramide (GlcCer) to ceramide and glucose. Gaucher disease (GD; OMIM #230800, 23090, and 23100) is an autosomal recessively inherited lysosomal storage disorder caused by biallelic pathologic variants in *GBA1* that result in deficient GCase activity and lysosomal accumulation of lipid substrates, including GlcCer and glucosylsphingosine (GluSph) [1]. In 2009, a multicenter international collaborative study conclusively established the connection between *GBA1* mutations and Parkinson disease (PD) highlighting *GBA1* variants as common genetic risk factors for the development of PD [2]. *GBA1* mutations also elevate the risk for dementia with Lewy bodies (DLB), providing further insight into the role of GCase in PD-related synucleinopathies [3]. This strong association spurred an upsurge in research on the cellular mechanism underlying the *GBA1*-PD link, as well as the potential of GCase as a therapeutic target for the synucleinopathies [4–6]. With the intensifying focus on GCase, the demand for reliable GCase antibodies for various applications has surged. In 2019, Qi et al evaluated several existing GCase antibodies for western blotting, revealing that many had limited utility [7]. While both academic and commercial groups have pursued the development of new GCase antibodies, a comprehensive characterization of these newer GCase antibodies for the research community has not been available. The development of GCase antibodies recognizing external epitopes essential for immunostaining, immunoprecipitation, and enzyme-linked immunosorbent assays (ELISA) has proven challenging. Here, we report the development of two mouse monoclonal antibodies, hGCase-1/17 and hGCase-1/23, raised against human GCase (hGCase) using the recombinant enzyme imiglucerase as the antigen. Characterization of these antibodies with genetic models, including *GBA1* loss-of-function human neuroglioma H4 line and human neurons differentiated from human embryonic stem cell (hESCs), demonstrated their superior specificity and potency in immunofluorescence, immunoprecipitation, and AlphaLISA (Amplified Luminescent Proximity Homogeneous Assay) assays.

These well-characterized hGCase antibodies will prove invaluable for future translational research and biomarker development in the context of GD, PD, and related synucleinopathies.

## Results

### Development of Monoclonal Antibodies Against Human β-glucocerebrosidase

Mice were immunized with imiglucerase (Cerezyme, Sanofi), the recombinant hGCase produced in Chinese hamster ovary cells for enzyme replacement therapy (ERT) in patients with GD [8]. Among the screened hybridoma clones, hGCase-1/17 and hGCase-1/23 exhibited selectivity for hGCase in initial screening assays including western blotting and immunofluorescence in HEK293-F cells overexpressing hGCase (Fig. S1). Notably, hGCase-1/23 demonstrated superior potency compared to hGCase-1/17 in both western blot and immunofluorescence assays. Neither of these antibodies cross-reacted with mouse β-glucocerebrosidase (mGCase) overexpressed in HEK293-F cells, underscoring their specificity for hGCase over mGCase.

### Generation of GCase Loss-of-function Human Neuroglioma H4 Cells

To support further hGCase-1/17 and hGCase-1/23 characterization, a GCase loss-of-function (*GBA1*^−/−^) human neuroglioma H4 line was generated. H4 (ATCC HTB-148) is a hypertriploid cell line with the modal chromosome number ranging from 63 to 78. To disrupt the multiple *GBA1* alleles in the H4 cell genome, zinc-finger nuclease (ZFN)-mediated cleavage was combined with targeted integration of a gene disruption cassette into exon 4 of *GBA1* alleles, following the workflow developed previously [9]. This targeting approach introduced a puromycin resistance cassette into the *GBA1* alleles (Fig. 1A). H4 cells with the integrated resistance cassette were enriched using puromycin selection, followed by single cell cloning. Established single cell clones were screened for the resistance cassette by PCR and subsequently with an *in vitro* GCase activity assay using fluorogenic substrate resorufin-β-D-glucopyranoside (Res-Glc). The most promising clone underwent Sanger sequencing, confirming frameshift indels in all *GBA1* alleles. A pair of PCR primers, P1 and P2, were designed to generate an amplicon of 761 bp, encompassing *GBA1* exons 4 and 5 for Sanger sequencing*. GBA1* allele(s) with the integrated puromycin resistance cassette could not produce PCR product due to the cassette’s size. The Sanger sequencing chromatogram of the chosen clone displayed double peaks near the Zinc-finger nuclease targeting site (Fig. 1B). Parsing the chromatogram with Poly Peak Parser (http://yost.genetics.utah.edu/software.ph) [10] indicated frameshift indels in the two *GBA1* alleles that were targeted by ZFN, but without the resistance cassette integration (Fig. S2A). Given the disruption of all *GBA1* alleles, this H4 clone is referred to as *GBA1*−/−, or the GCase knockout (KO) H4 line. *GBA1*-/- H4 cells lacked GCase activity in the Res-Glc *in vitro* GCase activity assay (Fig. 1C). Corroborating the absence of GCase activity, *GBA1*-/- H4 cells showed substantial GlcSph accumulation (Fig. 1D). Moreover, there were only minor changes in the GlcSph stereoisomer galactosylsphingosine (GalSph) in *GBA1*-/- H4 cells, indicating specific *GBA1* targeting (Fig. 1F).

**Figure 1.**
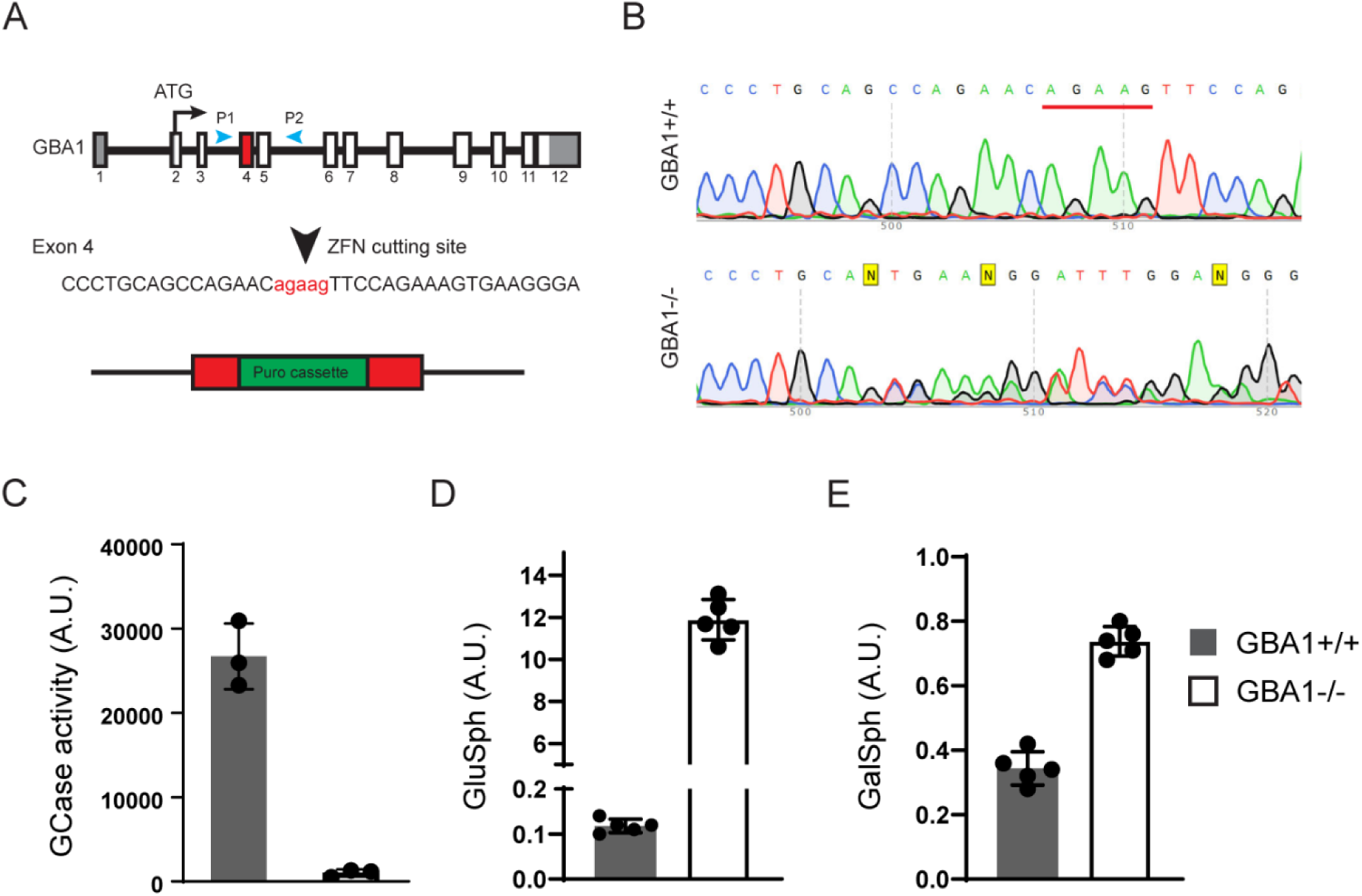
Generation of a GCase loss-of-function (*GBA1*−/−) human neuroglioma H4 line. (A) Disruption of *GBA1* alleles by targeting exon 4 with Zinc-finger nuclease (ZFN) and inserting a puromycin resistance cassette. Puromycin selection assisted with the enrichment of edited cells, which was followed by single cell cloning and validation by Sanger sequencing. (B) Sanger sequencing chromatogram showing double peaks in the region near the ZFN targeting site, demonstrating the presence of frameshift indel mutations in two *GBA1* alleles targeted by ZFN but lacking resistance cassette integration. (C) Lack of GCase activity in *GBA1*−/− H4 cells as determined with the Res-Glc *in vitro* GCase activity assay (*n*=3 biological replicates). (D, E) Glucosylsphingosine and galactosylsphingosine levels in the *GBA1+*/+ and *GBA1*−/− H4 cells (*n*=5 biological replicates).

### Benchmarking hGCase Antibodies in Traditional and Automated Capillary Western Blot

HGCase-1/17 and hGCase-1/23 were first evaluated by traditional western blotting using *GBA1*+/+ *and GBA1*-/- H4 cell lysates. This assessment compared these new antibodies with three recent commercial hGCase antibodies not yet independently evaluated: mouse monoclonal IgG 812201 (R&D Systems, MAB7410), mouse monoclonal IgG 2E2 (Abcam, ab55080), rabbit monoclonal IgG EPR5143(3) (Abcam, ab128879), and the well-characterized in-house antibody rabbit polyclonal IgG R386 [11] (Table 1). While 2E2, EPR5143, and R386 detected specific hGCase bands in *GBA1*+/+, but not GBA-/-, cells in traditional western blots, 812201, hGCase-1/17, and hGCase-1/23 yielded unsatisfactory results despite extensive optimization (Fig. 2A and data not shown). Subsequently, hGCase antibodies were compared in an automated capillary western blotting system (Simple Western from ProteinSimple, Bio-Techne), which automates the protein separation and immunodetection steps in traditional western blot, thus minimizing error-prone steps. The three commercial antibodies and R386 successfully detected hGCase via automated capillary western blotting, indicating their compatibility with this approach (Fig. 2B). Notably, 812201 produced better results in capillary western blotting than in traditional westerns. However, hGCase-1/17 and hGCase-1/23 failed to detect endogenous hGCase in *GBA1*+/+ cells using the capillary western blotting system. Hence, the three commercial hGCase antibodies were better suited for detecting endogenous hGCase in both traditional and capillary western blot. In contrast, hGCase-1/17 and hGCase-1/23 exhibited limited utility in western blot, detecting only overexpressed hGCase in HEK293-F cells (Fig. S1) but not at endogenous levels (Fig. 2).

**Table 1.** Antibodies used to detect hGCase by western blotting.

| hGCase Antibody | Vendor | Catalog # | Host | Clonality | Compatible Assays |
| --- | --- | --- | --- | --- | --- |
| hGCase-1/17 | Roche | Available soon | Mouse | Monoclonal | AlphaLISA, ICC, IP |
| hGCase-1/23 | Roche | Available soon | Mouse | Monoclonal | AlphaLISA, ICC, IP |
| 812201 | R&D systems | MAB7410 | Mouse | Monoclonal | Capillary WB |
| 2E2 | Abnova | H00002629-M01 | Mouse | Monoclonal | Conventional and Capillary WB |
| EPR5143 | Abcam | ab128879 | Rabbit | Monoclonal | Conventional and Capillary WB |
| R386 | In-house | N/A | Rabbit | Polyclonal | Conventional and Capillary WB |

**Figure 2.**
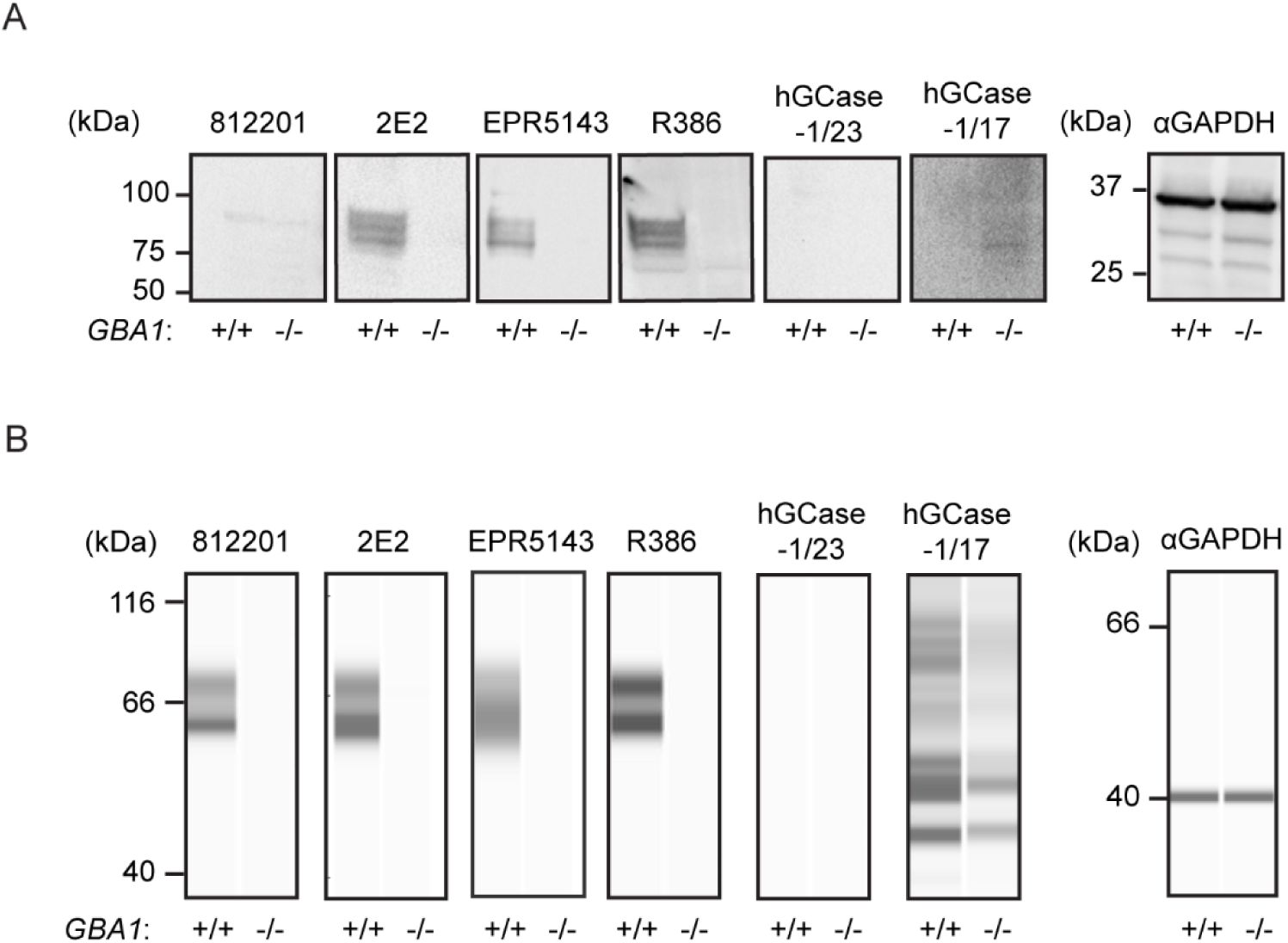
Performance of hGCase antibodies in detecting endogenous hGCase in traditional and automated capillary western blotting. A) Traditional western blotting analyzing *GBA1+*/+ and *GBA1*−/− H4 cell lysates with hGCase antibodies 812201, 2E2, EPR5143, R386, hGCase-1/17 and hGCase-1/23. B) Automated capillary western blotting analyzing *GBA1+*/+ and *GBA1*−/− H4 cell lysates with hGCase antibodies. Note that 812201 detected endogenous hGCase in *GBA1+*/+ H4 lysate in automated capillary western blotting.

### Characterization of hGCase-1/17 and hGCase-1/23 Through Immunostaining

Among the three commercial hGCase antibodies, 2E2 and 812201 were marketed as suitable for immunostaining assay. However, independent evaluation of 2E2 through immunostaining using *GBA1*+/+ and *GBA1*-/- SH-SY5Y cells yielded unsatisfactory results [12]. Consequently, hGCase-1/17 and hGCase-1/23 were compared with 812201 in immunostaining with paraformaldehyde (PFA)-fixed *GBA1*+/+ and *GBA1*-/- H4 cells. Both hGCase-1/17 and hGCase-1/23 displayed substantial selectivity towards endogenous hGCase, with staining confined to vesicular and tubular lysosomes that co-stained with the lysosomal membrane protein LAMP1, correlating with the expected lysosomal localization of hGCase (Fig. 3A and 3B). Impressively, *GBA1*-/- H4 cells exhibited virtually no background. Although both antibodies were selective towards hGCase, hGCase-1/17 exhibited weaker staining compared to hGCase-1/23, indicating a lower potency (Fig. 3B). Additional assessments of hGCase-1/23 in human neurons differentiated from *GBA1*+/+ and *GBA1*-/- human embryonic stem cell (hESC) lines further affirmed its specificity and potency (Fig. 3C). In contrast, 812201 failed to show specific staining in *GBA1*+/+ H4 cells (Fig. S3). Therefore, hGCase-1/23 emerged as the recommended choice for immunofluorescence assays.

**Figure 3.**
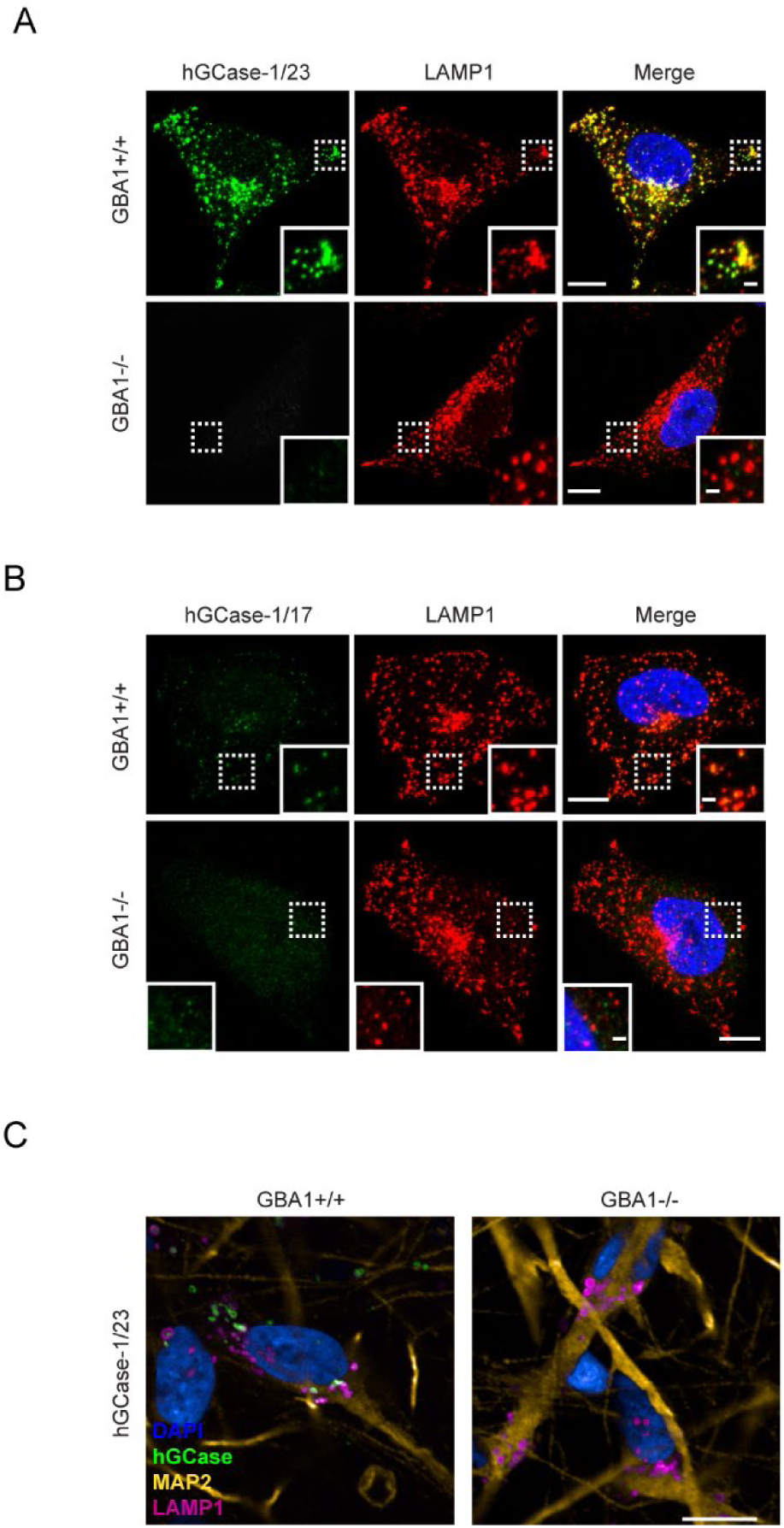
Specific immunostaining of hGCase in H4 cells and human neurons with hGCase-1/23 and hGCase-1/17. (A, B) *GBA1+*/+ and *GBA1*−/− H4 cells were fixed with 4% PFA, permeabilized with 0.05% Saponin, and stained with hGCase-1/23 (A) and hGCase-1/17 (B), together with LAMP1 antibody. Scale bars: 10 µm, and 1.5 µm in inserts. (C) *GBA1+*/+ and *GBA1*−/− human neurons were stained with hGCase-1/23. Scale bars: 25 µm.

### Immunoprecipitation of hGCase Using hGCase-1/17 and hGCase-1/23

Capitalizing on their success in recognizing external epitopes of hGCase in immunostaining, hGCase- 1/17 and hGCase-1/23 were tested in immunoprecipitation assays for their ability to capture hGCase in cell lysates. Both antibodies effectively pulled down hGCase in *GBA1*+/+ H4 cell lysates (Fig. 4A). Notably, hGCase-1/23 demonstrated superior capture efficiency than hGCase-1/17, in line with its superiority in the immunostaining assay (Fig. 4A). To further demonstrate their value in studying proteins interacting with hGCase, hGCase in human neuron lysates was captured with hGCase-1/23. Then, co-immunoprecipitated proteins were examined for the presence of lysosomal integral membrane protein-2 (LIMP-2/SCARB2), the trafficking receptor for hGCase [13–15]. LIMP-2 was co-immunoprecipitated with hGCase by hGCase-1/23 in *GBA1*+/+ but not *GBA1*-/- neuron lysates, indicating LIMP-2 co-immunoprecipitation relied on its interaction with endogenous hGCase (Fig. 4B).

**Figure 4.**
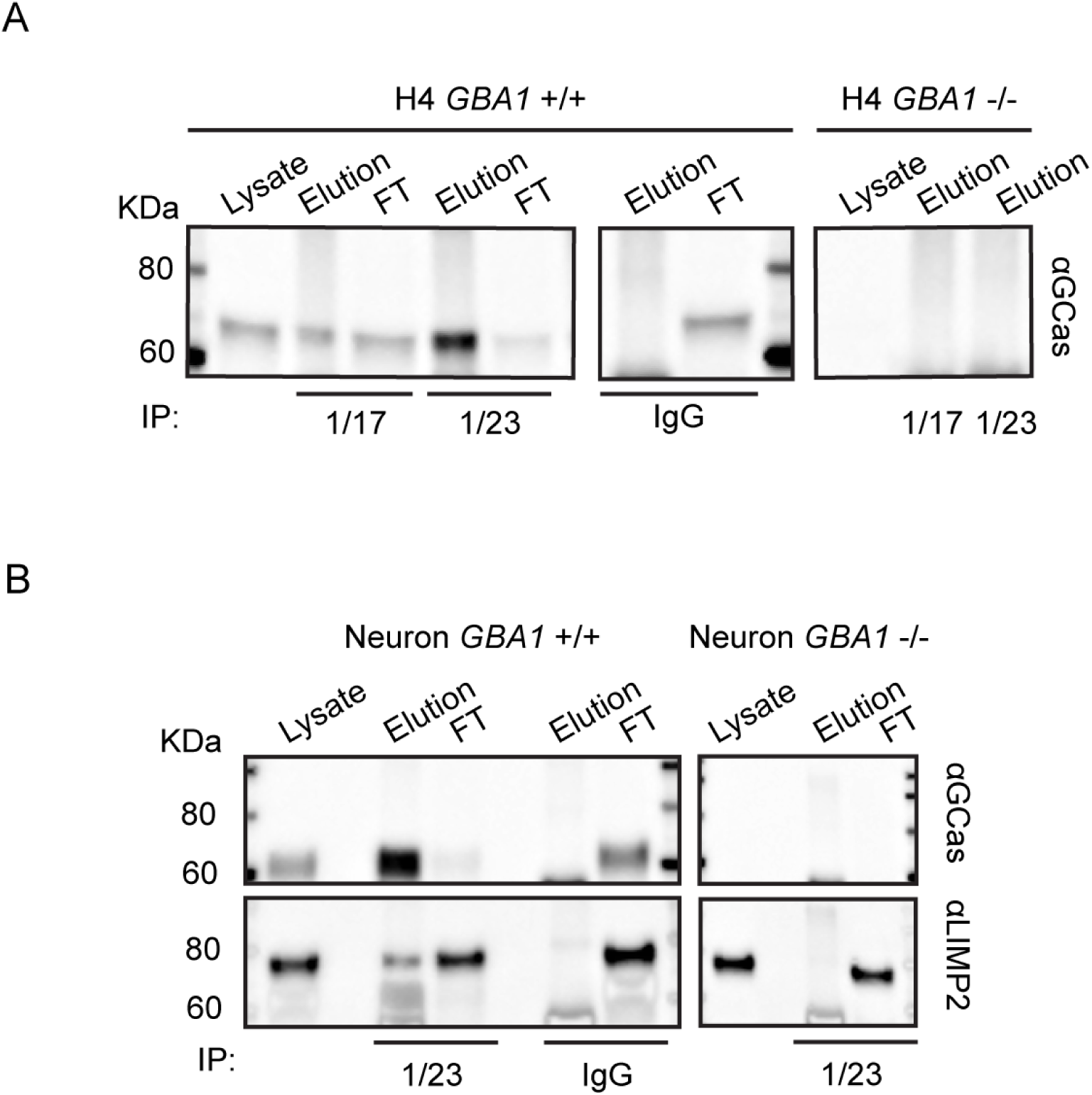
Application of hGCase-1/17 and hGCase-1/23 in immunoprecipitation. (A) HGCase in H4 cell lysates was immunocaptured with hGCase-1/17 or hGCase-1/23 and immunoblotted with hGCase antibody EPR5143 after elution. (B) Co-immunoprecipitation of LIMP2 with hGCase by hGCase-1/23. HGCase in neuron lysates was immunocaptured with hGCase-1/23, and the presence of LIMP2 in GCase co-precipitates was detected with a LIMP2 antibody.

### hGCase AlphaLISA Assay with hGCase-1/17 and hGCase-1/23

The AlphaLISA (Amplified Luminescent Proximity Homogeneous Assay) is a bead-based luminescent amplification assay developed by PerkinElmer to detect and quantify biomolecules in diverse sample types (Fig. 5A). It offers a streamlined workflow, enhanced sensitivity, expansive dynamic range, and adaptability for high-throughput applications. In the AlphaLISA assay, a standard sandwich configuration employs two distinct antibodies recognizing non-overlapping epitopes on the target molecule. One biotinylated antibody is coupled with an Alpha streptavidin-coated Donor bead, while the other antibody is directly conjugated to the AlphaLISA Acceptor bead. When the analyte is present, the Donor and Acceptor beads come together. Upon excitation with a 680 nm laser, a photosensitizer inside the Donor bead converts ambient oxygen to an excited singlet state. Singlet oxygen diffuses up to 200 nm to produce a chemiluminescent reaction in the Acceptor bead, emitting light at 615 nm proportional to the analyte amount.

**Figure 5.**
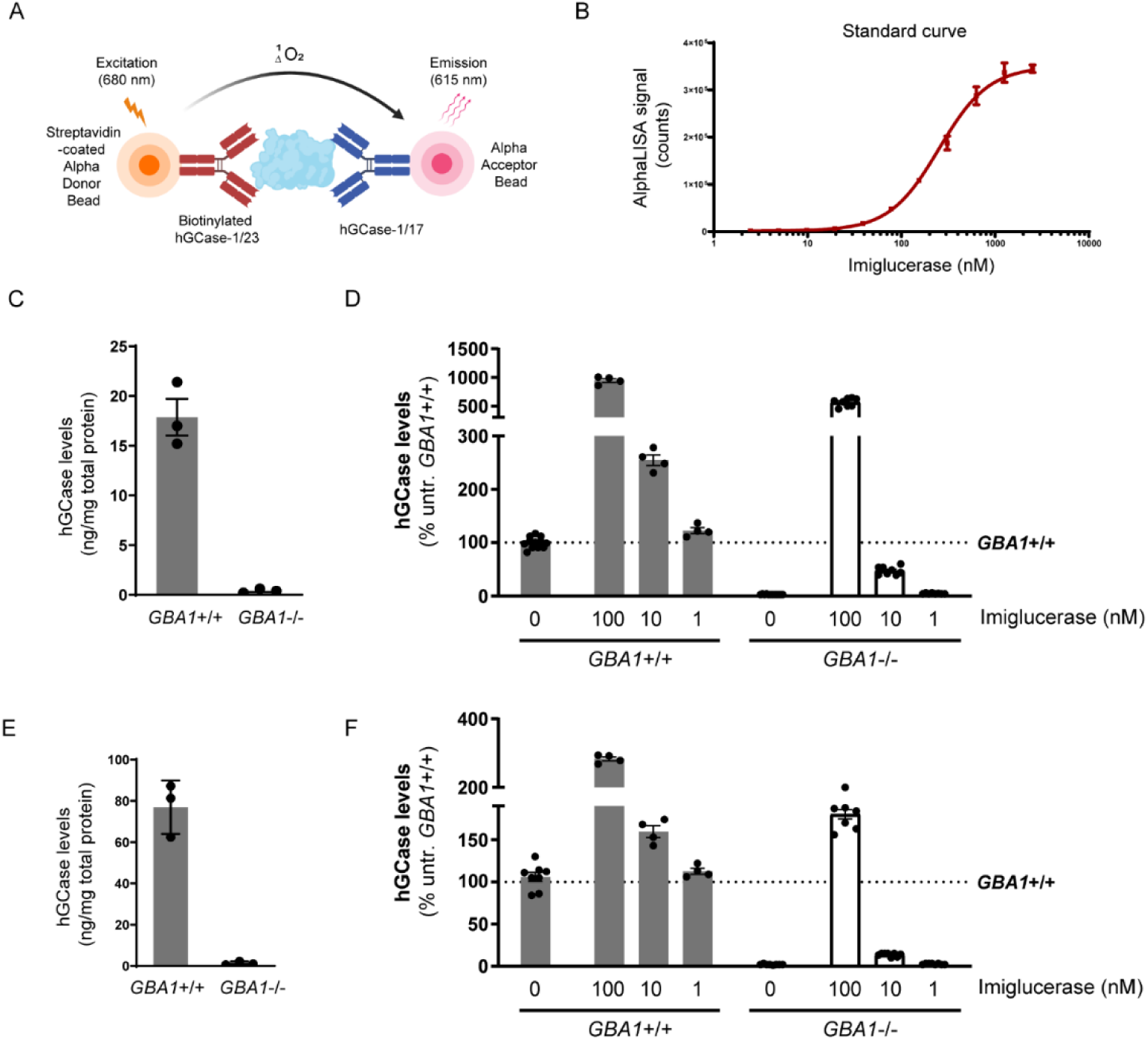
HGCase AlphaLISA assay. (A) In the sandwich AlphaLISA assay, hGCase-1/23 was biotinylated and associated with the Alpha Donor Bead, and hGCase-1/17 was directly conjugated to the Alpha Acceptor Bead. (B) Sensitivity and dynamic range of hGCase AlphaLISA assay (*n*=3 technical replicates). (C, D) Determination of hGCase levels in *GBA1*+/+ and *GBA1-*/- H4 cells with hGCase AlphaLISA assay. A 2 h treatment with imiglucerase led to a dose-dependent increase in hGCase levels (C, *n*=3 biological replicates; D, *n*=4 to 12 technical replicates). (E, F) Determination of hGCase levels in *GBA1*+/+ and *GBA1-*/- human neurons with hGCase AlphaLISA assay. A 2 h treatment with imiglucerase led to a dose-dependent increase in hGCase levels (C, *n*=3 biological replicates; F, *n*=4 to 12 technical replicates).

The potential of hGCase-1/17 and hGCase-1/23 for the AlphaLISA assay was explored by generating a dose-response curve with imiglucerase. The AlphaLISA assay utilized a sandwich configuration where hGCase-1/23 was biotinylated and associated with the Alpha Donor Bead, and hGCase-1/17 was directly conjugated to the Alpha Acceptor Bead. This assay effectively quantified imiglucerase levels, exhibiting exceptional sensitivity, broad dynamic range, and substantial signal-to-background ratio (Fig. 5B). With this assay, hGCase protein per milligram of cell lysate was estimated at approximately 20 ng in *GBA1*+/+ H4 cells and around 80 ng in *GBA1*+/+ neurons, while *GBA1*-/- H4 or human neuron cell lysates showed no hGCase protein, underscoring the assay’s specificity towards hGCase (Fig. 5C and E). Imiglucerase treatment increased hGCase levels in H4 cells and human neurons, regardless of *GBA1* genotype (Fig. 5D and F). This quantitative immuno-based assay has been previously employed to determine hGCase levels in PD and DLB postmortem brains [16].

## Discussion

This study unveils the development and characterization of hGCase-1/17 and hGCase-1/23, two mouse monoclonal antibodies with great utility for hGCase immunofluorescence, immunoprecipitation, and AlphaLISA assays. This achievement was made possible by immunizing with imiglucerase (recombinant full length hGCase) instead of peptide fragments, and by screening for selectivity at each stage of hybridoma clone selection. The resulting antibodies exhibited remarkable selectivity for hGCase and potent performance in immunofluorescence, immunoprecipitation, and AlphaLISA assays.

In immunofluorescence assays, hGCase-1/17 and hGCase-1/23 readily detected endogenous hGCase in *GBA1*+/+ H4 cells and human neurons, while exhibiting minimal background staining in *GBA1*-/- cells (Fig. 3). In immunoprecipitation assays, these antibodies successfully captured endogenous hGCase in both H4 cells and neurons. The co-immunoprecipitation of LIMP2 with hGCase demonstrated the antibodies’ value for investigating potential hGCase interactors (Fig. 4). Importantly, hGCase-1/17 and hGCase-1/23 bind to distinct epitopes on hGCase surface, enabling the development of the hGCase AlphaLISA assay with a sandwich configuration. This assay displayed high sensitivity and a wide dynamic range (Fig. 5). It holds potential for measuring hGCase as a translational biomarker and has already been used to determine GCase protein levels in PD and DLB postmortem brains [16].

Numerous pathological variants in *GBA1* compromise the trafficking of hGCase from the ER to lysosomes [18–20], and decreased levels of lysosomal GCase has been proposed to contribute to PD risk [21,22]. Consequently, hGCase is the therapeutic target in various disease-modifying therapies currently under development for PD. Strategies to restore lysosomal GCase activity include gene therapy to deliver exogenous hGCase [23] and small molecule chaperones to facilitate proper folding and trafficking of hGCase mutants [5,24–27]. The development of hGCase-1/17 and hGCase-1/23 now provides essential assays for evaluating the effectiveness of these therapeutic approaches in delivering hGCase to lysosomes in future clinical trials.

## Acknowledgements

We would like to thank Jean-Philippe Carralot and Nicole Soder for support with antibody generation, Judith Rothe and Karlheinz Baumann for support with the generation of *GBA1*-/- cell models, and Iris Ruf for support with lipid measurements.

The work of E.S., Y.C. and T.J. was funded by the Intramural Research Programs of the National Human Genome Research Institute and the National Institutes of Health. Additionally, this research was supported in part by Aligning Science Across Parkinson’s [ASAP-000458] through the Michael J. Fox Foundation for Parkinson’s Research (MJFF). For the purpose of open access, the author has applied a CC-BY public copyright license to all Author Accepted Manuscripts arising from this submission

## Methods

### Cell culture

H4 cells were maintained in DMEM with GlutaMAX (Gibco, cat no: 10566016), supplemented with Sodium Pyruvate (Gibco, cat no: 11360070) and 10% Fetal Bovine Serum (R&D, cat no: S12450) in 6- well tissue culture plates. Cells were kept in a 37°C incubator at 5% CO_2_. A full media change was performed every 2-3 days and cells were split by dissociation with TrypLE Express Enzyme (Gibco, cat no: 12605010).

### H4 cell pellet preparation

Cells were washed in PBS, dissociated with TrypLE Express Enzyme, quenched with cell media, and spun down at 500xg for 5 minutes. Cells were resuspended in PBS, transferred to a microcentrifuge tube, re-pelleted at 500xg for 5 minutes, and PBS was discarded. Pellets were flash frozen in dry ice and stored at −80°C for subsequent experiments.

### GBA1-/- human neurons

A *GBA1*-/- human embryonic stem cell (hESCs) line was previously generated and characterized [9]. Neural precursor cells (NPCs) were generated from the hESCs using a modified dual SMAD inhibition protocol [28], as previously described [29,30]. For neuronal differentiation, NPC were dissociated with 0.05% trypsin-EDTA (ThermoFisher, Cat. 25300054) and plated on poly-L-ornithine/laminin (PLO)-coated dishes at a density of 10,000–15,000 cells/cm2 in N2B27 medium supplemented with 100 ng/ml FGF-8 (PeproTech, Cat.100-25), 200 ng/ml sonic hedgehog (PeproTech, Cat. 100-45), and 100 μM ascorbic acid 2-phosphate (Sigma, Cat. A8960-5G) for seven days. Finally, the NPCs were plated on PLO-coated plates at a density of 50,000 cells/cm2 in BGAA medium, which was composed of N2B27 medium supplemented with 20 ng/ml BDNF (R&D System, Cat. 248-BDB-01M/CF), 10 ng/ml GDNF (PeproTech, Cat. 450-10), 500 μM Dibutyryl-cAMP (STEMCELL Technologies, Cat. 100-0244), and 100 μM ascorbic acid 2-phosphate (Sigma, Cat. A8960-5G). Neuronal cultures were maintained in BGAA medium for 6weeks, and the cell culture medium was changed every 4 days.

### Sanger sequencing

Genomic DNA was extracted from H4 cell pellets with Machery-Nagel genomic DNA extraction kit according to manufacturer instructions (cat no: 740952.250). Primers flanking exon 4 and exon 5 of *GBA1* were ordered from IDT. Sequence for forward primer (P1) is aaggcaggtctcaaactcctcac (5’ to 3’) and for reverse primer (P2) is agaatgggcagagtgagattctgc (5’ to 3’). The PCR mix was prepared by combining Ampligold Taq Mastermix (cat no: 439888) without GC enhancer with 0.5 pM of each primer, 0.5 ug of genomic DNA. PCR was programmed to the following conditions: 95°C 10 min for 1 cycle; 95°C 30 sec, 57°C 1 min, 72°C 1 min for 40 cycles; 72°C 7 min for 1 cycle and held at 4°C. A single specific band corresponding to the 761 bp amplicon was observed following DNA electrophoresis. PCR purification was performed with the QIAquick PCR Purification Kit (Qiagen, cat no: 28106) according to provided protocol. Sanger sequencing was prepared according to Genewiz Sample Submission guidelines using primers P1 or P2 mixed with PCR purification product. Sequencing results were analyzed using SnapGene software.

### Protein lysate preparation for Western Blot

150-200 µL of 1% Triton-X Lysis buffer [1% Triton X-100, 10% glycerol, 150 mM NaCl, 25 mM HEPES pH 7.4, 1 mM EDTA, 1.5 mM MgCl_2_, supplemented with one EDTA-free protease inhibitor tablet per 10 mL of buffer (Pierce, cat no A32965)] was added to H4 pellets from three confluent wells of a 6-well plate. Pellets were pulse-sonicated at 420 Hz for 10 quick pulses and rested for 15-30 minutes on ice. Then, the cells were centrifuged for 15 minutes at 13000 xg at 4°C and the supernatant containing lysate was collected in a new tube. Protein concentration was determined by Pierce BCA Protein Assay kit (ThermoScientific, cat no: 23225) and lysates were stored in −80°C.

### Resorufin β-D-glucopyranoside GCase Assay

5×10^4^ H4 cells were washed once with PBS and lysed in 30 µl GCase lysis buffer (0.05 M citric acid, 0.05 M KH2PO4, 0.05 M K2HPO4, 0.11 M KCl, 0.01 M NaCl, 0.001 M MgCl2, pH 6.0 with 0.1% (v/v) TritonX-100, supplemented with freshly added protease inhibitor). 10 µl of cell lysate were mixed with 10 µl of 10 mM resorufin-β-D-glucopyranoside and baseline fluorescence was measured at t_0_ immediately. The build-up of fluorescent product (resorufin) was measured after incubation for 2 h at 37 °C (*λ_ex_* = 535 nm and *λ_em_* = 595 nm) indicating GCase activity. Data was normalized to total protein levels and depicted as a percentage of untreated *GBA1* WT cells.

### Liquid chromatography-mass spectrometry analysis of GluSph

The following analytes and internal standards were purchased from Avanti Polar Lipids:

D-glucosyl-β-1-1’-D-erythro-sphingosine (GluSph (d18:1); #860535) and D-glucosyl-β-1-1’-D-erythro-sphingosine-d5 as internal standard 1 (GluSph-d5 (d18:1); #860636); D-galactosyl-β-1-1’-D-erythro-sphingosine (GalSph (d18:1); #860537) and D-galactosyl-β-1-1’-D-erythro-sphingosine-d5 as internal standard 2 (GalSph-d5 (d18:1); #860637). For chromatography HPLC grade solvents as well as Millipore water was used. Acetonitrile (LiChrosolv #1.00030) and methanol (LiChrosolv #1.06007) were obtained from Supelco (Merck), ammonium acetate for mass spectrometry was purchased by Sigma-Aldrich (#73594).

Stock solutions for analytes and internal standards were prepared at 1 mM in DMSO and kept at −20 °C. For further spiking solutions acetonitrile/water 9/1 (v/v) was used as solvent. Calibration solutions were prepared by serial dilution in acetonitrile/water 9/1 (v/v) containing 2% DMSO. The concentration ranged from C1 = 10 µM to C9 = 0.0039 µM.

Analysis was conducted on a LC-MS-MS system consisting of a Waters Xevo-TQ-S mass spectrometer connected to a complete Waters Acquity I-class UPLC system with a flow through needle sample manager using a mixture of acetonitrile/methanol/water 40/40/20 (v/v/v) as wash solvent. The auto- sampler temperature was set to 15 °C. The Xevo TQ-S instrument operated in positive ion electrospray mode with both quadrupoles tuned to unit mass resolution using nitrogen as nebulization- and desolvation gas. The nebulizer gas flow was set to 150 l/h and the desolvation gas flow to 800 l/h with a temperature of 500 °C. Argon was used as collision gas at a flow rate of 0.15ml/min. Analytes and internal standards were detected by multiple reaction monitoring mode (MRM) following the transitions m/z 462.3 to 282.3 and m/z 467.3 > 287.3 at a cone voltage of 30 V and a collision energy of 18 V.

Samples were analysed on a BEH glycan amide column (100 x 2.1 mm, 1.7 µm particle size, purchased from Waters, Switzerland) with a flow rate of 0.25 ml/min and an oven temperature of 30 °C. Eluent A consisted of 100mM ammonium acetate and for eluent B acetonitrile was used. Glycospecific separation was achieved by isocratic elution with 90% B followed by a washing step with 10% B and column reconditioning. The overall analysis time was 12 min.

### Sample preparation for *in vitro* GluSph measurements

50 µl of distilled water were added to deep frozen cell pellets and samples were shaken for 30 min at 25 °C until the cells thawed. Then, 100 µl methanol containing 0.005 µM internal standards was added. After shaking 5 min., the supernatant was transferred to a 96-well plate. Tubes were rinsed again with 100 µl of methanol and supernatant was transferred to the same plate. The samples were evaporated to dryness and reconstituted in 100 µl acetonitrile/water 90/10 (v/v) containing 1% DMSO and analyzed by LC-MS/MS. Lipids were quantified using the peak area ratio analyte/internal standard (=response).

### Antibody Details

For antibody details, please see Table 1.

### Western blot: Conventional chemiluminescence platform

For each sample, 40 µg of protein was loaded onto a 4–20% Mini-PROTEAN TGX PVDF gel (Bio-Rad Laboratories). After transfer with the Trans-Blot Turbo transfer system (Bio-Rad Laboratories), PVDF membranes (Bio-Rad Laboratories) were dried and reactivated with methanol. Methanol was rinsed off with water and TBST [1xTBS (from 10x TBS, 0.1% Tween] and the membrane was blocked for 1 hour at RT with blocking solution [1×TBST, 5% (w/v) BSA (Sigma, cat no: A-7888)]. 5% (w/v) NFDM (Biorad, cat no: 1706404) blocking was also performed for each antibody; 5% BSA showed clearer bands and less background compared to 5% NFDM blocking for all antibodies. PVDF membranes were probed overnight at 4°C with primary antibodies in antibody diluent [2% (w/v) BSA, 1x TBST]. Primary antibodies were diluted so that each antibody had a final concentration of approximately 0.5 µg/mL. PVDF membranes were washed 3 × 10 minutes at RT with TBST. This was followed by incubation with Goat HRP-coupled secondary anti-mouse IgG (Abcam, cat no: ab205719, 1:10K dilution) or anti-rabbit IgG (Abcam, cat no: ab 205718, 1:60K dilution) antibodies in antibody diluent. PVDF membranes were washed 3 × 10 minutes with TBST. The antigen–antibody complexes were detected with SuperSignal West Pico PLUS Chemiluminescent Substrate (ThermoScientific) using the ChemiDoc XRS+ Imager System (Bio-Rad). Blots were exposed for 3 seconds, and images were generated every 6 seconds.

### Western blot: Automated capillary platform

Simple Western’s Fluorescence Separation Module kit and 12-230 kDa Fluorescence Separation Capillary Cartridges (cat no: SM-W004) were used to perform automated western blotting. Samples were prepared and loaded according to manufacturer instructions. Samples were prepared in 0.1x sample buffer (10x Sample Buffer diluted in water) and Fluorescence 5x Master Mix such that 5 µg of protein was loaded per lane at a concentration of 1 mg/mL. Prepared samples in master mix as well as Biotinylated Ladder were boiled at 95°C for 10 minutes. All primary antibodies were diluted in the provided Antibody Diluent 2 at 1:50 dilution, except for R386 and GAPDH, which were diluted 1:1500 in Antibody Diluent 2. 10 µL of diluted antibody was loaded in each lane. 10 µL of HRP-conjugated mouse or rabbit secondary antibody was loaded to the corresponding lanes. Finally, 15 µL peroxide-luminol mix was loaded into each lane, the cartridge was centrifuged for 5 min at 1000xg, and wash buffer was added according to the user manual. The cartridge and capillaries were loaded into the Jess instrument (ProteinSimple) and the loading times used were the default for HRP chemiluminescence detection. Results were visualized and analyzed with the Compass for Simple Western program.

### Immunocytochemistry

Cells were seeded in 8-well imaging chambers at a density of 30K cells/chamber and grown for 48 hours with daily feeding. After 48 hours, cells were washed once in PBS and fixed in 4% paraformaldehyde (20 min, RT). Fixed cells were washed in PBS three times and incubated in primary antibody ICC antibody diluent [0.1% Saponin (filtered) and 1% BSA in PBS] overnight at 4°C. Wells were washed three times with PBS, followed by 3 × 5 min washes with PBS, then covered with aluminum foil and incubated with secondary antibody diluted in ICC antibody diluent in RT. Three PBS wash and 3 × 5 min washes in PBS were repeated, and cells were stored in PBS with NucBlue Fixed Cell ReadyProbes Reagent (DAPI) (ThermoFisher, cat no: R37606) in 4°C until imaging. Confocal images were acquired using a Zeiss LSM 880 confocal microscope using the 63x oil objective.

### AlphaLISA

One million H4 cells were washed once with PBS and lysed in 80 µl GCase lysis buffer (0.05 M citric acid, 0.05 M KH2PO4, 0.05 M K2HPO4, 0.11 M KCl, 0.01 M NaCl, 0.001 M MgCl2, pH 6.0 with 0.1% (v/v) TritonX-100, supplemented with freshly added protease inhibitor). Samples were diluted 1/5 or 1/10 in 1X Immunoassay buffer (Perkin Elmer, #AL000F). A 2-fold serial dilution of imiglucerase (Genzyme) ranging from 1250 pM − 2.4 pM was generated as a standard curve. 10 µl of samples or standards were incubated for 4 hours at RT in the dark with 1 nM of biotinylated hGCase-1/23 and 20µg/ml of hGCase-1/17- conjugated Acceptor beads (beads:antibody 50:1). Subsequently, 40 µg/ml of AlphaScreen Streptavidin Donor beads (Perkin Elmer, #6760002) were added and incubated 1 hour at RT in the dark. Plate was read on a Tecan Spark plate reader with ex: 680 nm and em: 520-620 nm. GCase concentration was calculated based on a sigmoidal non-linear regression of the imiglucerase standard and then normalized to total protein concentration.

### Immunoprecipitation

For immunoprecipitation, the Dynabeads™ Protein A Immunoprecipitation Kit (Thermo Fisher, #10006D) was used. 10 µg of hGCase-1/17 or hGCase-1/23 or mouse IgG (Merck Millipore, #NI03) were coupled to 50 µl of Protein A magnetic Dynabeads in 200 µl Ab Binding and Washing Buffer for 30 minutes at RT on a rotating wheel. Subsequently, supernatant was removed, and Ab-conjugated beads were mixed with 100 µg of cell lysate and incubated for 1 hour at RT on a rotating wheel (20 µg of cell lysate were saved for analysis). After incubation, supernatant (= flowthrough, FT) was saved for analysis and beads were washed 3 times in 200 µl Washing Buffer. For elution, beads were resuspended in 20 µl of Elution Buffer mixed with 7 µl of 4X LDS sample buffer (Thermo Fisher, # NP0007) and 3 µl of 10X reducing agent (Thermo Fisher, # NP0004) and heated for 10 min to 70 °C. Supernatant (= elution) was saved for analysis. Samples were analysed by SDS-PAGE loading 10 µl of each fraction per lane (lysate and FT were mixed with appropriate volumes of 4X LDS sample buffer and 10X reducing agent prior to loading). Proteins were transferred to a nitrocellulose membrane using the iBlot 2 device and membranes were blocked for 1 hour at RT in 5% (w/v) Blotting Grade Blocker/PBS (Biorad, #170-6404). GCase was detected using rb mAb to hGBA antibody (abcam, #128879, clone EPR5143(3)) at 1/1000 in blocking solution. After 3 x 5 minutes washing in PBS, membranes were incubated with goat anti-rabbit HRP-labelled antibody (Perkin Elmer, # NEF812001EA) at 1/10000 in 1 % (w/v) Blotting Grade Blocker/PBS for 1.5 hours at RT. Proteins were detected using the SuperSignal West Dura Extended Duration Substrate Kit (Thermo Fisher, #34076).

## Supplementary Figures

**Figure S1.**
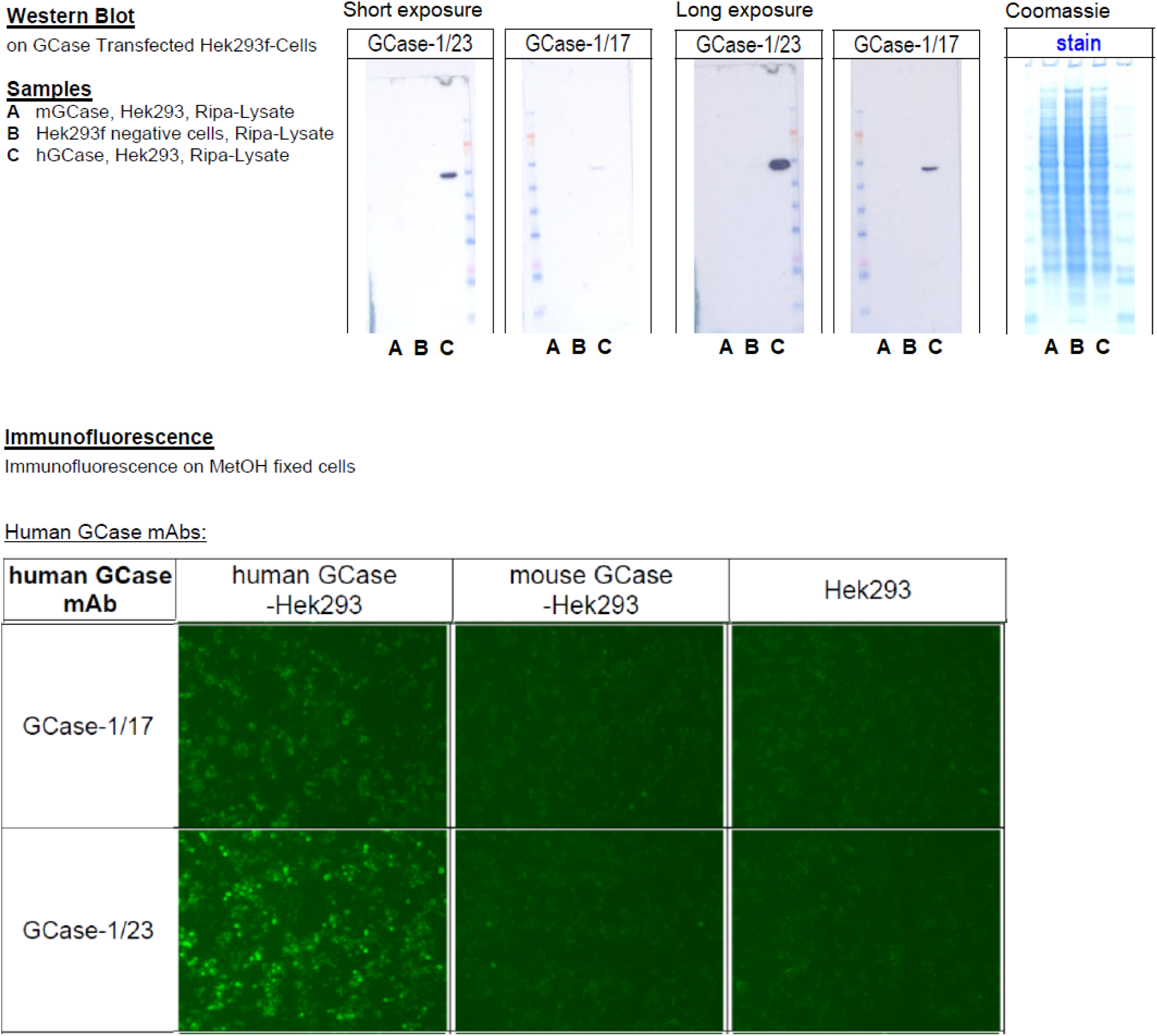
Characterization of hGCase-1/17 and hGCase-1/23 hybridoma clones. Hybridoma clones hGCase-1/17 and hGCase-1/23 were derived from mice immunized with imiglucerase. HGCase-1/23 demonstrated stronger potency than hGCase-1/17 towards hGCase overexpressed in HEK293-F cells in western blotting and in immunofluorescence assays. Neither of the two antibodies cross-reacted with mouse β-glucocerebrosidase (mGCase) overexpressed in HEK293-F cells, demonstrating their species specificity toward hGCase.

**Figure S2.**
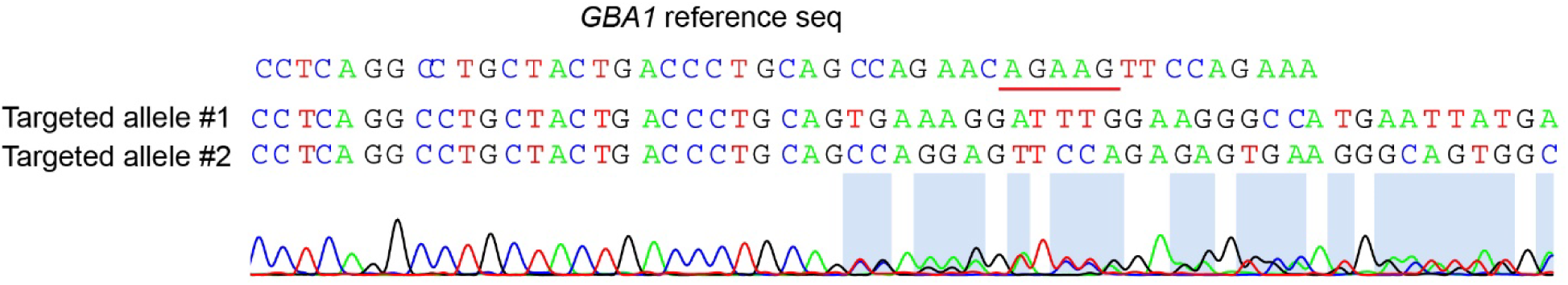
Frameshift indels in the two *GBA1* alleles targeted by ZFN but lacking resistance cassette integration. Sanger sequencing *GBA1* alleles without resistance cassette integration showed double peaks in the chromatogram indicating the two *GBA1* alleles were targeted by ZFN as well. Parsing the double peak chromatogram using Poly Peak Parser revealed frameshift indels in both alleles.

**Figure S3.**
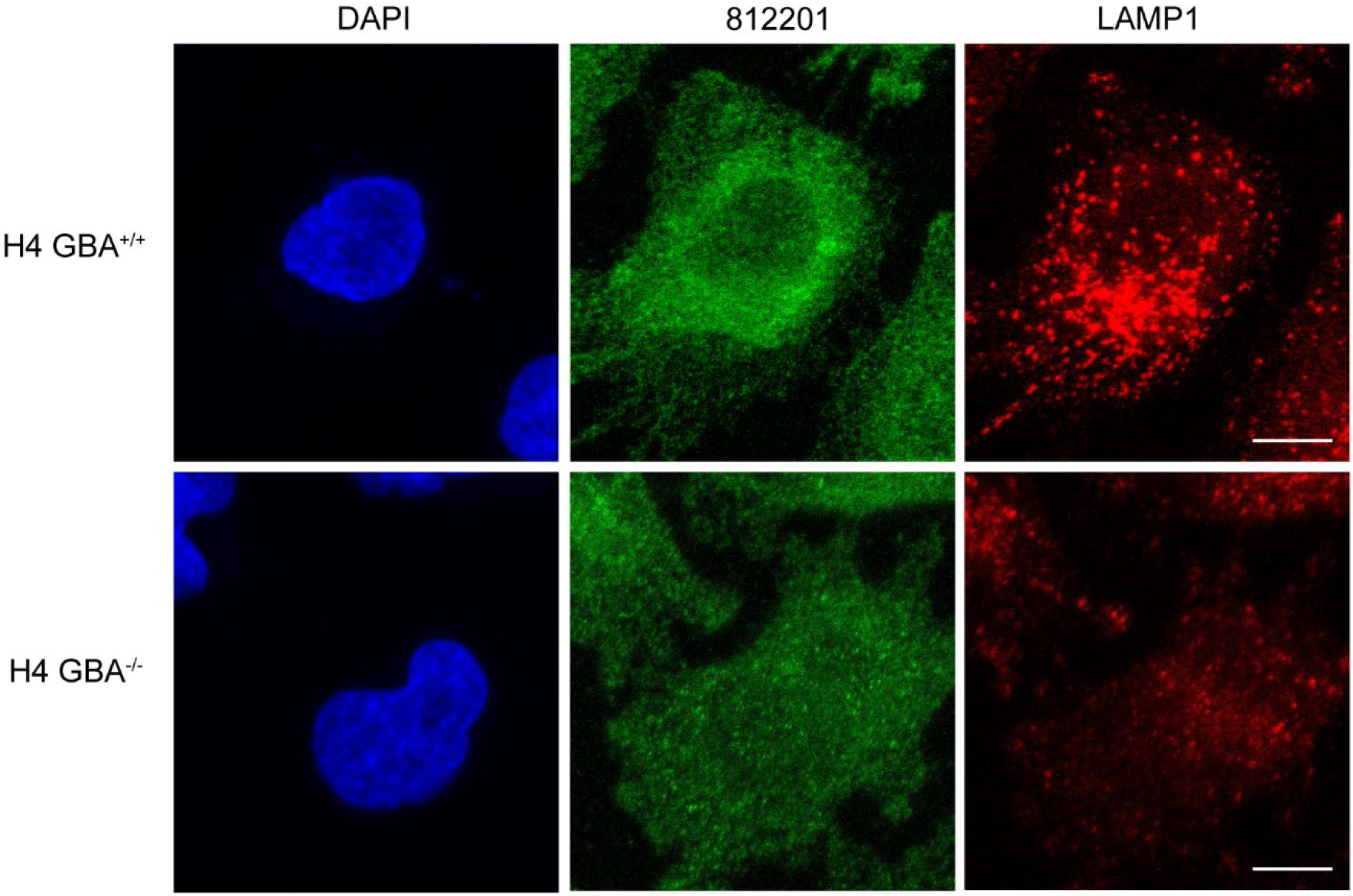
No specific immunostaining of hGCase in H4 cells with hGCase antibody 812201. *GBA1+*/+ H4 cells were fixed with 4% PFA, permeabilized with 0.05% Saponin, and stained with hGCase antibody 812201, together with LAMP1 antibody. The staining pattern was diffusive showing no localization in lysosomes marked with LAMP1. Scale bar: 10 µm.

